# Resolution of subcomponents of synaptic release from post-synaptic currents in rat hair-cell/auditory-nerve fiber synapses

**DOI:** 10.1101/2020.05.13.070920

**Authors:** Eric D. Young, Jingjing Sherry Wu, Mamiko Niwa, Elisabeth Glowatzki

**Affiliations:** Center for Hearing and Balance; Department of Biomedical Engineering; Department of Otolaryngology-Head and Neck Surgery; Department of Neuroscience; Center for Sensory Biology, Johns Hopkins School of Medicine, Baltimore MD 21205 USA

## Abstract

The synapse between inner hair cells and auditory nerve fiber dendrites shows large EPSCs, which are either monophasic or multiphasic. Multiquantal or uniquantal flickering release have been proposed to underlie the unusual multiphasic waveforms. Here the nature of multiphasic waveforms is analyzed using EPSCs recorded *in vitro* in rat afferent dendrites. Spontaneous EPSCs were deconvolved into a sum of presumed release events with monophasic EPSC waveforms. Results include: first, the charge of EPSCs is about the same for multiphasic versus monophasic EPSCs. Second, EPSC amplitudes decline with the number of release events per EPSC. Third, there is no evidence of a mini-EPSC. Most results can be accounted for by versions of either uniquantal or multiquantal release. However, serial neurotransmitter release in multiphasic EPSCs shows properties that are not fully explained by either model, especially that the amplitudes of individual release events is established at the beginning of a multiphasic EPSC, constraining possible models of vesicle release.

## INTRODUCTION

The synapse between an inner hair cell (IHC) and the dendrite of an auditory-nerve fiber (ANF) is specialized to serve the needs of auditory encoding, which requires high rates of discharge and precise timing of neural spikes evoked by acoustic signals (Matthews and Fuchs 2010; Rutherford and Moser 2016). In the mature mammalian cochlea, spike timing in ANFs is determined by the timing of single EPSCs, and thus by the temporal properties of neurotransmitter release from the IHC (Siegel 1992; Yi et al. 2010; Rutherford et al. 2012; Wu et al. 2016). An understanding of the physiology of this synapse is an essential part of a comprehensive description of auditory encoding.

The IHC has a synaptic ribbon which is thought to organize vesicles for high-rate release (Frank et al. 2010; Buran et al. 2010; Becker et al. 2018; Jean et al. 2018). In electron-microscopic examinations, there are multiple vesicles that are apparently in position for immediate release under the ribbon (Matthews and Fuchs 2010; Frank et al. 2010; Chakrabarti et al. 2018) and others associated with the ribbon away from the membrane. Examination of postsynaptic EPSCs in mammalian ANF dendrites has revealed a range of waveforms. EPSCs are either monophasic, with a smooth rise to a single peak followed by a near-exponential decay, or multiphasic, with multiple peaks (Grant et al. 2010). The latter likely are responses to multiple rapid sequential releases of neurotransmitter (Glowatzki and Fuchs 2002; Keen and Hudspeth 2006; Chapochnikov et al. 2014). This range of EPSC waveforms raises the question of the nature of vesicle release, in particular whether it is uniquantal (one synaptic vesicle per EPSC) or multiquantal (Neef et al. 2007; Grant et al. 2010; Chapochnikov et al. 2014; Grabner and Moser 2018). Clearly release is multiquantal in some hair-cell preparations (e.g. frog, Li et al. 2009, 2014; turtle, Schnee et al 2013) and in similar ribbon synapses in rat retinal bipolar cell (Singer et al. 2004), but the extent of multiquantal release in mammalian cochlear hair cells is still unclear. Mechanisms by which multiphasic EPSCs could arise from either the release of a single vesicle or the asynchronous release of several vesicles have been proposed (Glowatzki and Fuchs 2002; Matthews and Fuchs 2010; Chapochnikov et al. 2014; Vincent et al. 2018).

Chapochnikov and colleagues (2014) argued for uniquantal release in which the multiphasic EPSCs are produced by “flickering”, rapid opening and closing, of the fusion channel during the release of one vesicle. This conclusion was based on theoretical arguments supporting the adequacy of uniquantal release to produce the observed EPSCs and on data consistent with univesicular release, including capacitance recordings in IHCs (Grabner and Moser 2018) and measurement of the charge injected by EPSCs, which showed that multiphasic EPSCs inject the same charge into the postsynaptic terminal as monophasic ones (Grant et al 2010; Chapochnikov et al 2014). While this evidence is compelling, it is based on a relatively small number of recordings.

Here we further analyze the question of the nature of EPSCs using a large database of recordings of spontaneous EPSCs in afferent dendrites of hearing rats. Deconvolution is used to separate the EPSCs into a sum of elementary monophasic events which are assumed to correspond to successive releases of neurotransmitter. From 50% to 90% of EPSCs are monophasic with symmetrical, single-mode amplitude and charge distributions, the rest are multiphasic. The charge injected by monophasic and multiphasic EPSCs is roughly the same, consistent with previous findings (Grant et al. 2010; Chapochnikov et al. 2014), showing that vesicle release is from pools of neurotransmitter of fixed size, such as a constant number of vesicles (one or more.. The properties of EPSC amplitudes are also consistent with the fixed-pool model. However, analysis of the details of serial neurotransmitter release in multiphasic EPSCs show properties that are not fully explained by either uniquantal or multiquantal models.

## RESULTS

EPSCs were recorded in the apical turn of isolated rat (P17-P31) cochleas, *in vitro.* We used data from patch voltage-clamp recordings on the dendritic terminals of ANFs contacting IHCs; the experiments were done at room temperature. To maximize synaptic currents, the dendritic terminals were hyperpolarized by voltage clamp (−84 to −99 mV). EPSCs occurred without explicit stimulation, except for depolarizing the IHC membrane potential by 5.8 (3 recordings) or 15 (5 recordings) mM potassium in the extracellular solution. In 5.8 mM K^+^ the IHC membrane potential is close to rest (~ −60 mV) and in 15 mM K^+^, it is depolarized (~ −40 mV). In these data, EPSCs are adult-like in their properties, as described previously (Wu et al. 2016). Data from eight preparations (Table 1) containing between 1,384 and 19,362 EPSCs are included; the average rates of EPSCs ranged between 2 and 58 /s (Wu et al. 2016, Figure 3).

**Table 1.**
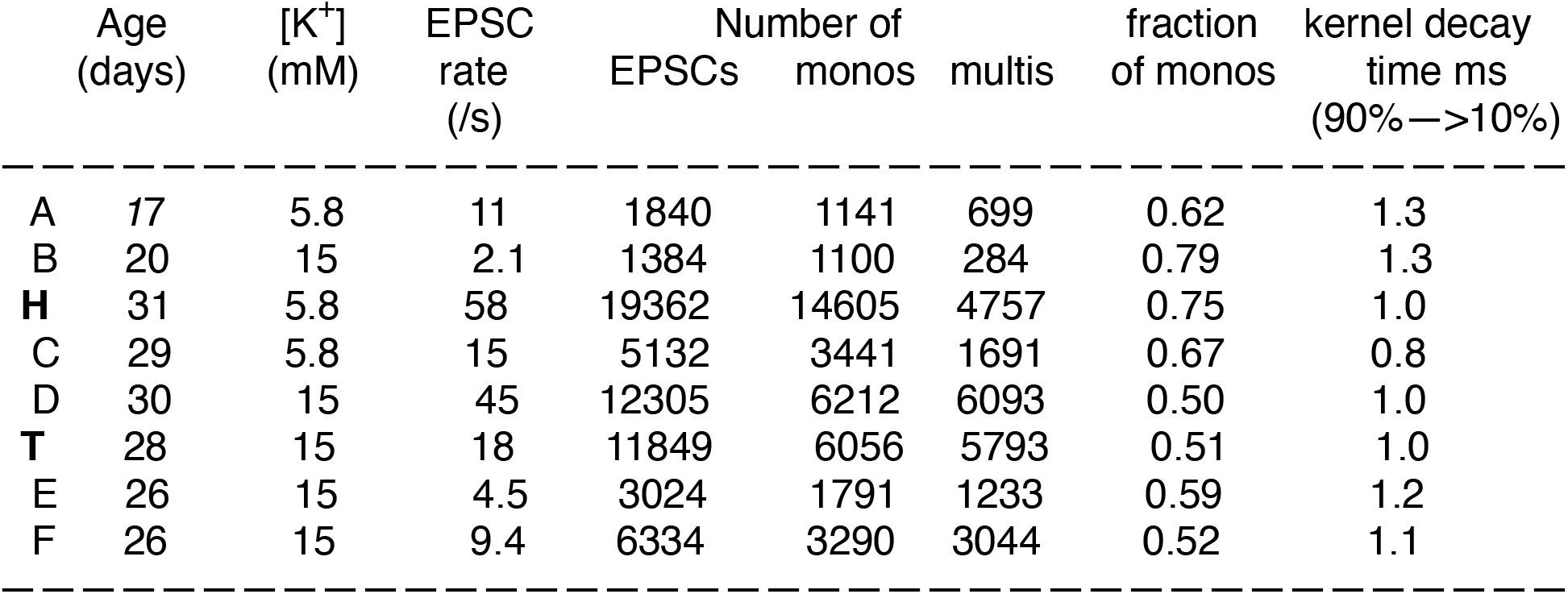
Summary of the 8 experiments. Total numbers of EPSCs are given, as well as the number of monophasic and multiphasic EPSCs. Kernel decay time is time from 90% to 10% of maximum value, as in Figure 7. The experiments are identified in the first column, using the figure numbers (A, B, …) from Figures 2 and 3 - figure supplement 1, along with H and T.

|  | Age<br>(days) | [K<sup>+</sup>]<br>(mM) | EPSC<br>rate<br>(/s) | Number of<br>EPSCs | monos | multis | fraction<br>of monos | kernel decay<br>time ms<br>(90%—>10%) |
| --- | --- | --- | --- | --- | --- | --- | --- | --- |
| A | 17 | 5.8 | 11 | 1840 | 1141 | 699 | 0.62 | 1.3 |
| B | 20 | 15 | 2.1 | 1384 | 1100 | 284 | 0.79 | 1.3 |
| H | 31 | 5.8 | 58 | 19362 | 14605 | 4757 | 0.75 | 1.0 |
| C | 29 | 5.8 | 15 | 5132 | 3441 | 1691 | 0.67 | 0.8 |
| D | 30 | 15 | 45 | 12305 | 6212 | 6093 | 0.50 | 1.0 |
| T | 28 | 15 | 18 | 11849 | 6056 | 5793 | 0.51 | 1.0 |
| E | 26 | 15 | 4.5 | 3024 | 1791 | 1233 | 0.59 | 1.2 |
| F | 26 | 15 | 9.4 | 6334 | 3290 | 3044 | 0.52 | 1.1 |

### Measuring EPSC waveforms

Examples of EPSCs are shown by the negative-going blue waveforms in Figure 1A. EPSCs in this preparation show a variety of amplitudes and shapes, as illustrated here (Glowatzki and Fuchs 2002; Grant et al. 2010). These range from “monophasic” waveforms with a single peak (the examples labelled “1”) to complex “multiphasic” ones (multipeak waveforms labelled with numbers greater than 1). The shapes of the multiphasic EPSCs suggest that they might represent a summation of monophasic waveforms occurring asynchronously over the duration of the EPSC. Under this hypothesis, the monophasic waveform represents the release of a single bolus of neurotransmitter.The bolus could be from all or part of one synaptic vesicle or from the simultaneous release of several vesicles. The analysis makes no a-priori assumptions about which.

**Figure 1.**
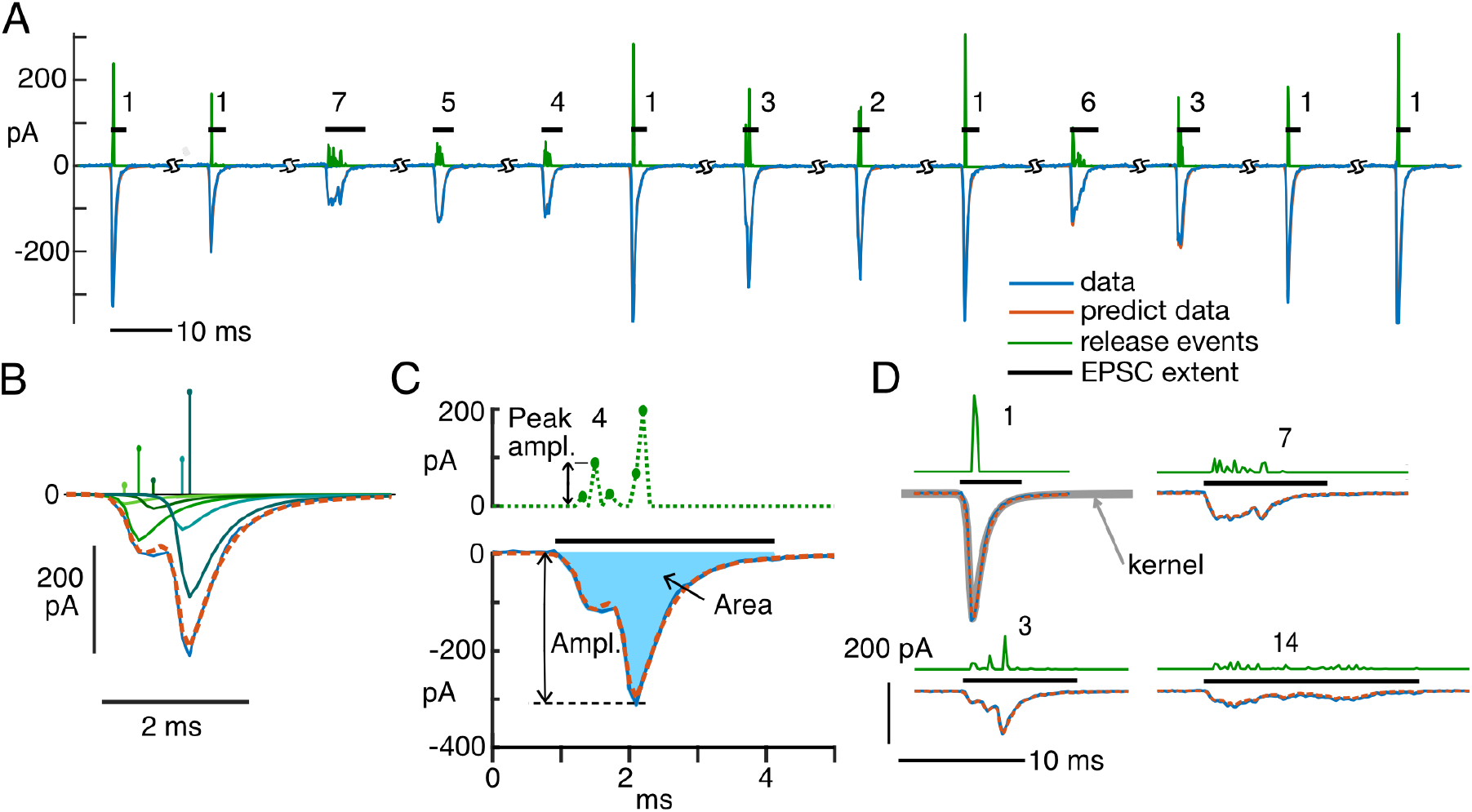
Analysis of monophasic and multiphasic EPSCs. A. 13 examples of EPSCs (blue negative waveforms), the sequence of kernels producing the best fits (whose amplitudes and times are shown by the green positive waveforms), and the fits calculated from the event sequences (red, mostly hidden behind blue). The EPSCs are shown in the order In which they occurred in one experiment, except that segments of baseline have been removed, where marked. The numbers are the numbers of events (defined in the text) in the fit of the EPSC. The horizontal black bars show the non-zero extent of each EPSC. B. Detailed view of one EPSC (blue negative trace) and the calculated fit (overlying dashed red trace), with the five kernels in the fit waveform plotted separately below to show how they sum to approximate the EPSC. The kernel amplitudes (*a_j_* in Eqn. 1) and times of occurrence (*t_j_*) are plotted as impulses above. The kernels are greenish in color, with hues coded to match the amplitudes. C. The same EPSC as in B. The peak amplitude of the EPSC (“Ampl.”) and the area (“Area”, filled with blue) are marked. The dashed green waveform at top is the event sequence as in B, except plotted as dots connected with dashed lines to emphasize that there are zero values in between most of the events. Immediately adjacent events are gathered together in classifying EPSCs, as described in the text. Thus there are 4 events in this fit, the last two non-zero kernels are combined into one event. D. Expanded view of four EPSCs with calculated event sequences and fits, labelled by the number of events in the fit. The kernel used for all parts of this figure is shown by the gray line overlying the monophasic EPSC (labelled “1”, top left). Note that the kernel closely overlies the waveform of this monophasic EPSC. In fact, the variance of the monophasic waveforms was very small; they generally varied only in amplitude.

The shape of the monophasic EPSC (here called the kernel) reflects all the dynamics of generating the EPSC, including neurotransmitter release, transport, binding, and channel gating. We assume that multiphasic EPSCs are the result of asynchronous release of differently-sized packets of neurotransmitter, each of which is subject to the same dynamics, producing the same monophasic waveforms; these monophasic waveforms summate linearly to produce the multiphasic waveforms (Eqn. 1, Figure 1B). To analyze these waveforms, deconvolution was used to estimate the set of monophasic waveforms (kernels) making up a multiphasic EPSC (Figure 1B; see Methods for details). The computation finds the set of kernels, in terms of their amplitudes and times of occurrence, that provides the best fit to the EPSC waveform, subject to the constraint that the smallest possible number of kernels should be used (lasso method, Tibshirani, 1996). The deconvolution results in a series of impulses (the green positive signals in Figure 1) located at the times of kernel occurrences (releases of boluses of neurotransmitter) with amplitudes equal to the amplitudes of the kernels (Figures 1C, D; see also Andor-Ardo et al. 2012; Chapochnikov et al. 2014).

For every experiment, a kernel is computed as the average of isolated, apparently monophasic EPSCs in the data record, meaning ones with short simple waveforms that are separated by at least 10 ms from other EPSCs. An example of a kernel waveform is plotted with a thick gray line in Figure 1D (the upper left plot labelled “1”), superimposed on a monophasic EPSC. Kernels from the 8 experiments are shown in Figure 1-figure supplement 1.

Figure 1B illustrates the deconvolution of an EPSC into 5 kernels (called “events”). The events are shown as impulses, vertical lines whose time and height are the kernel time and amplitude. The corresponding kernel waveforms are plotted below, with the EPSC waveform (blue) and the sum of the kernels (red dashed). Figure 1C shows the deconvolution result for the same EPSC as in Figure 1B, now with dashed lines connecting the kernel amplitudes and the sample points with no kernel in between. Generally, the kernels are isolated with zeros in between, as in the first three events in this example. However, there are often non-zero values at successive (adjacent) sample times (spaced by 0.1 ms here); in the analysis below, these are assumed to form a single event for the sake of classifying EPSCs (as in the 4th and 5th events in Figures 1C). Of course, how the events are parsed makes no difference to the calculation of the fit or its accuracy. The amplitude of an event is the sum of the kernel amplitudes contained within it.

The measures used to characterize EPSCs are shown in Figure 1C, bottom plot. EPSCs are characterized by: (1) the number of events in the EPSC, determined as described in the previous paragraph; (2) the EPSC amplitude (the current at the negative minimum of the EPSC); and (3) the area of the EPSC (the region shaded blue) multiplied by the sampling increment. This product is the total charge delivered to the terminal by the EPSC. We consider charge to be a more meaningful measure of the strength of the EPSC than the amplitude, which is highly variable, depending on waveform, for multiphasic EPSCs; in fact, the amplitude is shown below to vary with the degree of synchrony of the release events. The use of charge is supported by data from dual recordings showing that postsynaptic EPSC charge is proportional to the presynaptic capacitance change in a prolonged depolarization of the hair cell, a direct measure of the amount of neurotransmitter released (Li et al. 2009). Moreover, charge is a measure of the postsynaptic excitation that will be produced by the EPSC, because for brief currents, the EPSC charge flows mainly into the membrane capacitance, which has the effect of depolarizing the membrane by an amount proportional to the charge. Across the range of EPSC rates (Table 1), most EPSC properties are the same. This includes their amplitudes and areas, the fraction of monophasic versus multiphasic EPSCs, and the number of release events within multiphasic EPSCs. One property of EPSCs, the decay time, is weakly correlated with rate, shown in Figure 7C.

### EPSCs are mostly monophasic; multiphasic EPSCs are smaller in amplitude

The number of events in the best fit to an EPSC provides a way to characterize its qualitative nature (monophasic vs multiphasic for example). In Figures 1A and 1D, the green positive waveforms show the event sequence calculated as described above for that EPSC. The numbers next to each EPSC give the number of events in the signal. Note in Figure 1A that the largest-amplitude EPSCs are monophasic and the multiphasic EPSCs are smaller in amplitude, especially those with more than 4 events. This is expected, for example, if each EPSC represents the release of a fixed (except for random variation) total amount of neurotransmitter, so that the amplitude of a multiphasic EPSC is reduced by desynchronization in the release process.

EPSCs most commonly show monophasic waveforms, 50–79% in the eight experiments analyzed here (Table 1 and Figure 2C). In Figure 2C, data are shown separately (gray and dashed colored plots) for each experiment and as an average across all experiments (black). Note that the ordinate of this plot is scaled logarithmically in order to show the small numbers of EPSCs containing larger numbers of events. Although monophasic waveforms predominate, there is a surprisingly large number of multiphasic EPSCs, different from most synapses.

**Figure 2.**
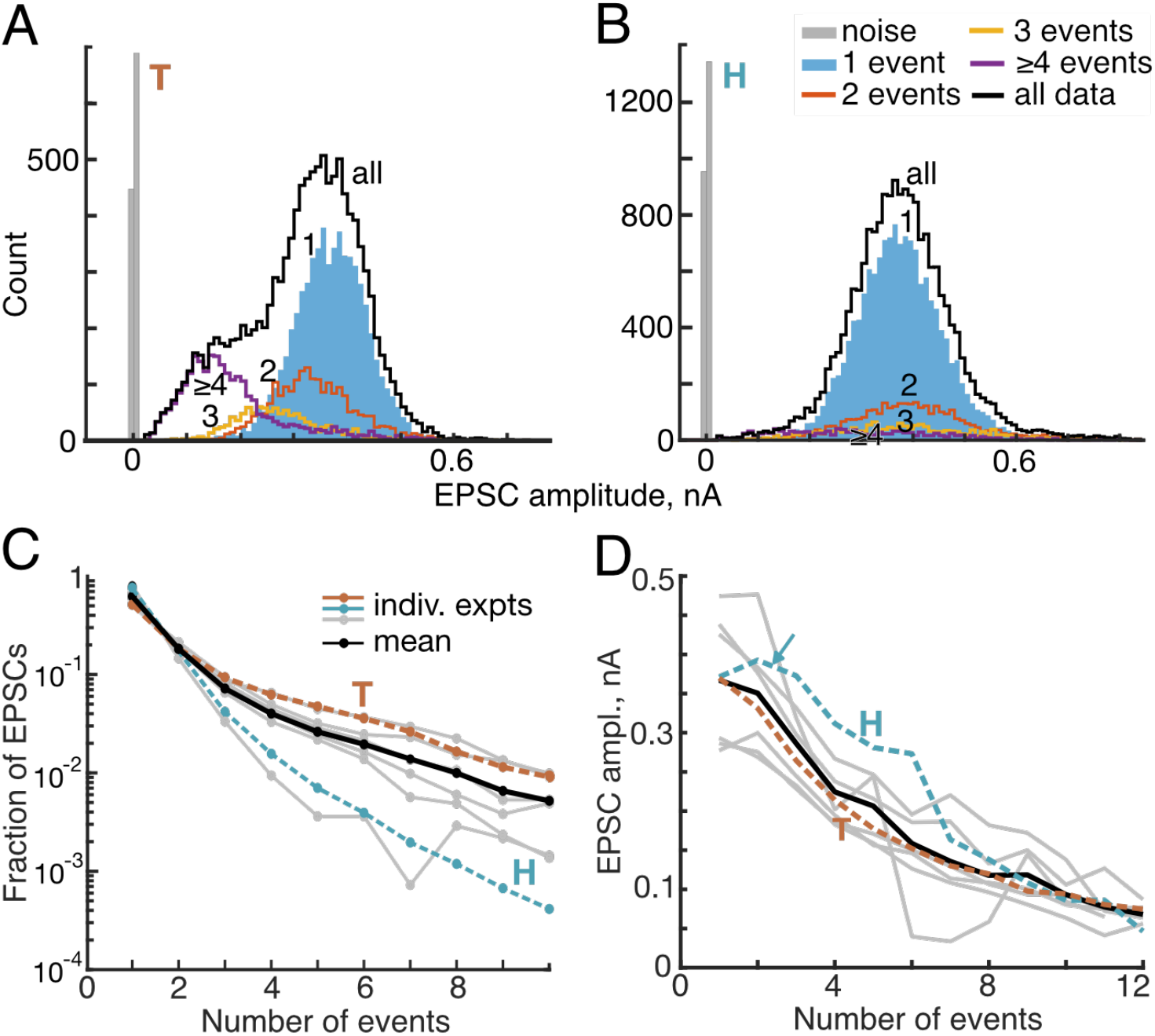
Amplitudes of EPSCs vary with the number of events in the EPSC waveforms. A,B. Histograms of EPSC amplitudes for data from two example experiments (“T” and “H”, identified in the same way in all figures). EPSCs with 1, 2, 3, and ≥4 events (i.e. combining all EPSCs with 4, 5, etc. events) are plotted separately (legend in B). The black histograms show all the data together and the gray histogram represents samples drawn at random from regions outside all EPSCs. In the data for experiment H, the 3 and ≥4 histograms are scaled up by 2x for clarity. C. The fraction of EPSCs with various numbers of events for eight experiments (gray and colored lines). The mean across all experiments is the black line. The dashed and colored lines show data from example experiments T (brown) and H (cyan). EPSCs with 1 event (monophasic) are most numerous, and the prevalence of EPSCs decreases rapidly and monotonically as the number of events increases. Note that the ordinate is logarithmic. D. Mean values of EPSC amplitude versus the number of events in the EPSC for eight experiments and the overall mean, plotted as in part C. The decrease is clear (R=−0.96, P<<0.001). The arrow points out an exception to the usual decrease in amplitudes for Experiment H. Experiment T was studied in 15 mM K^+^ and experiment H in 5.8 mM K^+^.

In this and subsequent figures, data from two experiments are identified with brown (experiment “T”) and cyan (experiment “H”) dashed lines. These two experiments were chosen because T is *typical* of the population in all measures, whereas H is an outlier in some measures. They are both excellent data sets with large numbers of EPSCs (11849 and 19362, respectively) from very stable recordings. Experiment H has the *highest* EPSC rate (58 /s) and also contains the second largest fraction of monophasic waveforms (75%), whereas T has a medium EPSC rate (18 /s) and contains 51% monophasic waveforms.

The distributions of EPSC amplitudes are shown for the two example experiments in Figures 2A and 2B for all EPSCs and EPSCs having 1, 2, 3, and at-least-4 events (see legend). The mean EPSC amplitude decreases with the number of events in the EPSC, seen clearly by the leftward shift of the distributions of experiment T. The same trend holds in the data from experiment H, but is less apparent because of the small number of multi-event EPSCs. Nevertheless, the mean amplitudes decline, usually monotonically, with the number of events in the EPSCs in all experiments (Figure 2D; R=−0.96, P<<0.001; Figure 2 - figure supplement 1). Note that mean amplitudes of EPSCs having 2 and 3 events are larger than the amplitudes of monophasic EPSCs for experiment H (cyan arrow). The reason for this is discussed in the next section.

The amplitude distributions in Figure 2A,B and Figure 2 - figure supplement 1 do not show evidence of mini-EPSCs. The traditional analyses of the amplitudes of EPSCs or EPSPs based on summation of various numbers of simultaneously released vesicles assume a monophasic waveform shape for a single-vesicle release (the mini-EPSC). However, the monophasic EPSCs (blue histograms labelled “1”) have a uniform symmetric Gaussian-like distribution of amplitudes with no small-amplitude peak. The overall amplitude distributions (black lines labelled “all”) may have a tail that extends toward zero amplitude (7/8 experiments, H being the exception). However, these low-side tails are always the multiphasic EPSCs, clearly shown by the T experiment in Figure 2A. In this experiment and five others (in Figure 2 - figure supplement 1), the only possible mini-EPSCs are multiphasic, but that is inconsistent with the usual models of synapse function.

### EPSC area is approximately constant, independent of event number

To test the hypothesis that EPSCs containing different numbers of events result from asynchronous release of neurotransmitter from a pool of fixed size (e.g. a single vesicle or a fixed number of vesicles), and not recruitment of different sized release pools (e.g. different numbers of vesicles), we examined the areas of EPSCs. The area is equal to the charge delivered by an EPSC and is a measure of the total neurotransmitter received by the synapse, assuming that no non-linear effects occur (discussed later in Figure 7). Thus, under the assumption of a fixed pool of transmitter, the area should be constant regardless of the number of events in an EPSC (Grant et al. 2010; Chapochnikov et al. 2014). This expectation is approximately realized in the data. Distributions of the EPSC area are shown in Figure 3A and 3B for the two example experiments. The areas are shown separately according to the number of events per EPSC (1, 2, 3, and ≥4, see the legend). In all eight experiments (Figures 3A,B and Figure 3 - figure supplement 1), the areas of monophasic EPSCs (the blue histograms labelled “1”) are symmetrically distributed with a single mode. The coefficient of variation (SD divided by the mean) of these histograms varies between 0.21 and 0.27 (median 0.24) for the eight experiments, consistent with usual results on the variability of EPSCs in central synapses (Matthews and Fuchs, 2010).

**Figure 3.**
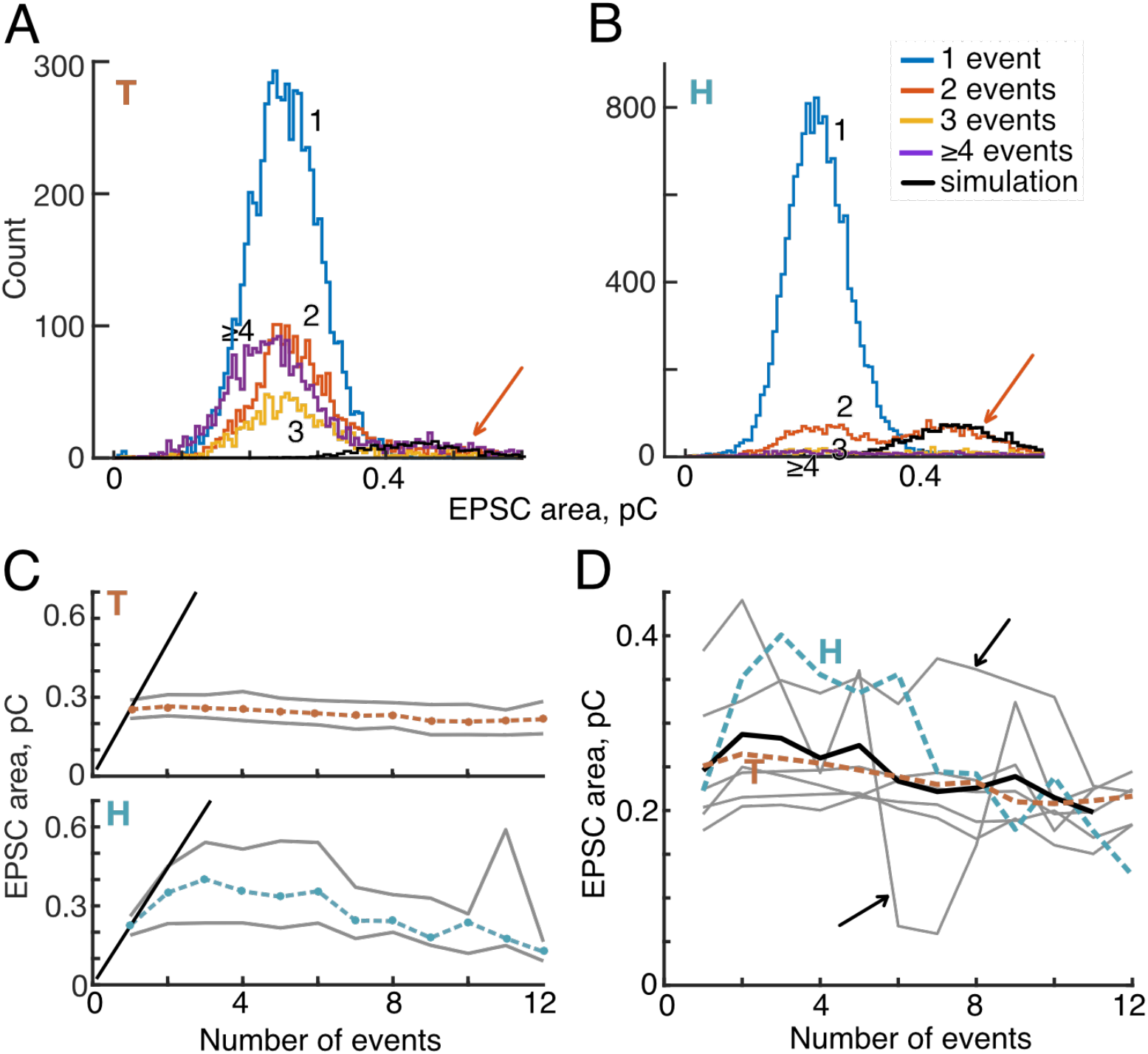
EPSC area declines slightly as the number of events in the EPSC increases. A. B. Histograms of EPSC areas for two example experiments. Most experiments are similar to example T in that areas show a roughly normal distribution which is similar for different numbers of events. The distributions deviate from normal by having truncated tails. In panel B, the histogram for 2-event EPSCs (orange) shows a noticeable second peak (orange arrow) at the expected place if these EPSCs represent overlap of EPSCs from two releases of neurotransmitter occurring close together (see text). The black lines show the tail distributions computed from the single-EPSC area distribution as described in text. T in panel A shows a smaller second peak. C. Plots of the mean area (dashed colored lines) plus and minus 1 SD (gray lines) versus the number of events in the EPSC. Data are shown for Experiments T and H. The diagonal black lines at left are the expected behavior if multiple-event EPSCs result entirely from overlapping 1-event EPSCs, for which area should increase linearly with the number of events. D. Mean EPSC area plotted versus event number, for eight experiments (gray and colored lines, plotted as in Figure 2D); the mean of the eight experiments is the black line. The data show small but significant declines in area as event count increases (R=−0.84, P=0.0006). Points are plotted only if they are based on at least 10 EPSCs. Most experiments (5/8) show the same behavior as experiment T (brown line). Two experiments (black arrows) have few data for 5 and more events, so are noisy.

Experiment T is typical of most of the data in that the distributions of areas are similar for various numbers of events; the areas scatter around roughly the same mean value. In fact, the mean values of area histograms usually decline slightly when plotted against the number of events, as for experiment T in Figure 3C (upper plot).

Often, the area histograms for multievent EPSCs have a right-side tail formed by EPSCs with areas larger than the 1-event EPSCs. Such a tail is clearly shown in Figure 3B by the 2-event histogram (orange) (for experiment H, at areas > 0.35, orange arrow); another example is shown in Figure 3 - figure supplement 1C. Such tails are largest in experiments, like H, with high rates of EPSC production. They occur when two EPSCs occur close together in time; if two EPSCs are sufficiently close, then the waveforms will partially merge and the algorithm for defining EPSCs will classify them as part of the same EPSC, with 2 or more events. The resulting composite EPSC has an area twice as large as isolated EPSCs; it may also have a larger amplitude, depending on the degree of overlap (e.g. the cyan arrow in Figure 2D for experiment H). To test this idea, the 2-event area histogram expected from such overlaps was calculated from the 1-event histogram and the inter-event (time) interval histogram (near-exponential as in Wu et al. 2016) and shown to correspond well to the upper tail of the observed 2-event histogram (the black lines in Figure 3B and Figure 3 - figure supplement 1C, see also Chapochnikov et al. 2014; Niwa et al. 2020 personal communication). Small upper tails are also seen in experiment T (Figure 3A).

The plot of mean area for experiment H in Figure 3C (bottom plot) shows clearly this effect of waveform overlap in that the mean area for 2- through 6-event EPSCs exceeds the mean for 1-event EPSCs substantially (by up to 1.8). The black diagonal lines at left in Figure 3C show the location of the data predicted if the area of EPSCs is proportional to the number of events in the waveform, the expected behavior if the larger areas reflect release from a larger neurotransmitter pool (e.g. as many vesicles as there are events in the EPSC). The data from experiment H lie well below the black diagonal line, consistent with the fact that the generation of such EPSCs results from a mixture of true 2-event releases and the overlap of adjacent 1-event EPSCs. The decrease of the area for larger numbers of events is expected because of the low likelihood of many EPSCs occurring close enough in time to overlap.

The difference in behavior between experiments H and T for 2-and 3-event EPSCs is striking. The T experiment also shows right-side tail distributions, but the number of EPSCs in those tails is small and has little effect on the mean areas. The mean area curves computed from eight experiments (Figure 3D) are mostly similar to the result for experiment T. There is a small but steady decline of area with event count. Two experiments had few EPSCs at larger numbers of events, and their plots are quite variable (black arrows), but still show a general decrease with event number. The overall mean across all 8 experiments (black line) shows the same trend, a small decrease of area with event number (but still significant R=−0.84, P=0.0006). Thus the hypothesis of neurotransmitter release from a fixed pool is supported by the data, although the small decrease in area for multiphasic EPSCs still needs explanation.

### Lack of serial dependence of the event numbers

Figure 1 shows an apparently random ordering of the numbers of events in subsequent EPSCs. However, serial order effects are often observed in hair cell and ANF activity (Lowen and Teich 1992; Peterson et al. 2014; Wu et al. 2016), and it is worthwhile to look for similar effects on the synchrony of release. There is not a serial dependence of the number of events in an EPSC, as measured by the conditional mean event-number in the n^th^ EPSC, given the event-number in the (n-1)^th^ EPSC (Figure 4A). The conditional mean event count (ordinate) is normalized by the overall mean count in the same experiment, to deemphasize differences in mean count across experiments. This lack of dependence was confirmed by computing the mutual information between the event numbers for successive pairs and triples of EPSCs; this measure is very small (median 0.02 bits, NS) meaning that there is no information in the previous one or two EPSCs about the number of events in an EPSC. There is also no effect of the amplitude or area of an EPSC on the event number in the following EPSC (not shown).

**Figure 4.**
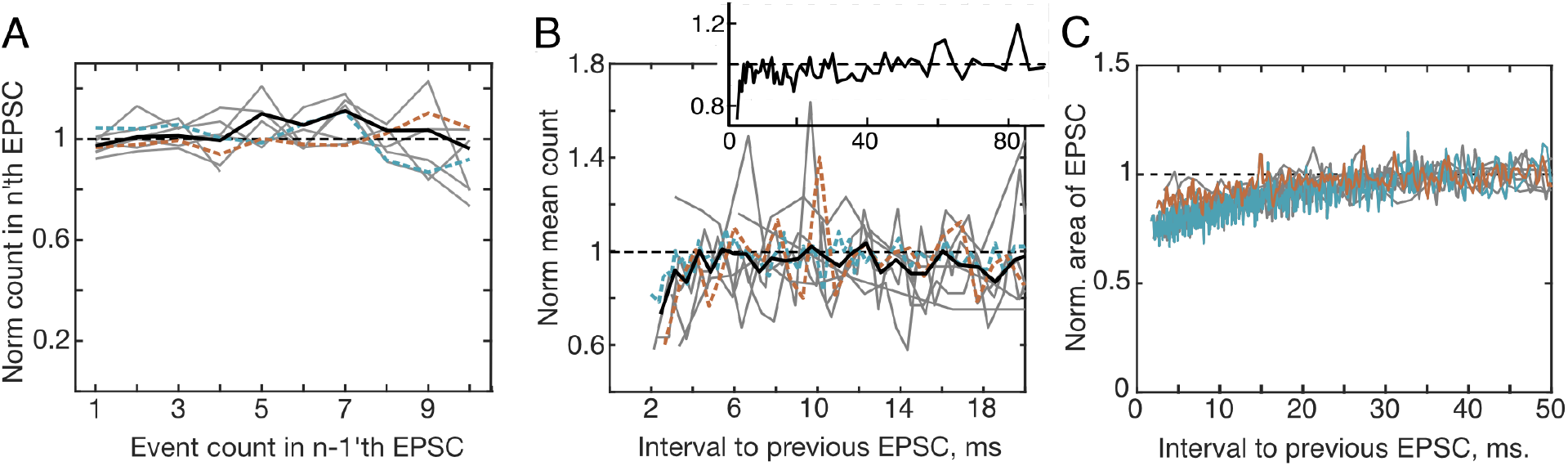
Serial dependence between event counts in successive EPSCs. A. Conditional mean counts in an EPSC plotted against the event count in the previous EPSC. Plots are shown for eight experiments (gray or colored, as in Figure 2D and 3D) and the overall mean (heavy black line); the counts are normalized by the overall mean count in the same experiment. B. Normalized mean event count plotted against the interval to the previous EPSC, for pairs with the first EPSC having only 1 event. The count is reduced slightly for EPSCs that follow a previous EPSC closely (< 6 ms). The normalization is done by dividing by the mean count of the pairs with inter-EPSC intervals >40 ms in the same experiment. The heavy black line is the mean across experiments and the inset shows the mean on a longer time axis. C. The area of EPSCs also decreases at short inter-EPSC interval, with a time constant slightly longer than that observed for event counts. Area is normalized by dividing by the mean area for pairs with inter-EPSC intervals >40 ms.

Two rather weak serial dependence phenomena are observed in these data. First, the number of events per EPSC is smaller for EPSCs occurring at short intervals (less than about 6 ms) following a previous EPSC (Figure 4B). EPSCs are longer in duration for larger event numbers, because of the asynchronous occurrence of neurotransmitter releases. Because this lengthening constrains the possible inter-event intervals, the analysis in Figure 4B was done using only pairs of EPSCs in which the first EPSC has one event, so is as short as possible. Repeating the analysis with all pairs gave the same result as in Figure 4B. The count is again normalized across experiments by dividing by the mean event counts for intervals >40 ms from the same experiment. The dip in event number at short intervals amounts to about 0.5 event on an absolute scale (without normalization). The recovery of event count appears to show two phases, a fast phase that lasts around 6 ms, shown in the main part of Figure 4B, and a slower phase that is not fully recovered until about 40-50 ms, shown by the inset which is the same mean data on a longer time scale. The second serial-dependence phenomenon is that the area or amplitude of EPSCs decreases slightly for EPSCs spaced closer than 20 ms (shown for area in Figure 4C).

### Event organization within EPSCs

If EPSCs containing different numbers of events draw from roughly constant-sized pools of neurotransmitter, as argued from Figure 3, it is interesting to ask how that pool is divided up among the events in the EPSC. This includes both the timing and the charge-contents of the events. Figure 5A shows ten examples of EPSCs (blue negative signals from experiment T) containing five events (green positive signals) and the calculated best-fitting model EPSCs (red, mostly occluded by the blue plots), plotted as in Figure 1C. There is a diversity of patterns of events, both in terms of amplitudes and occurrence times, which is typical of the data.

**Figure 5.**
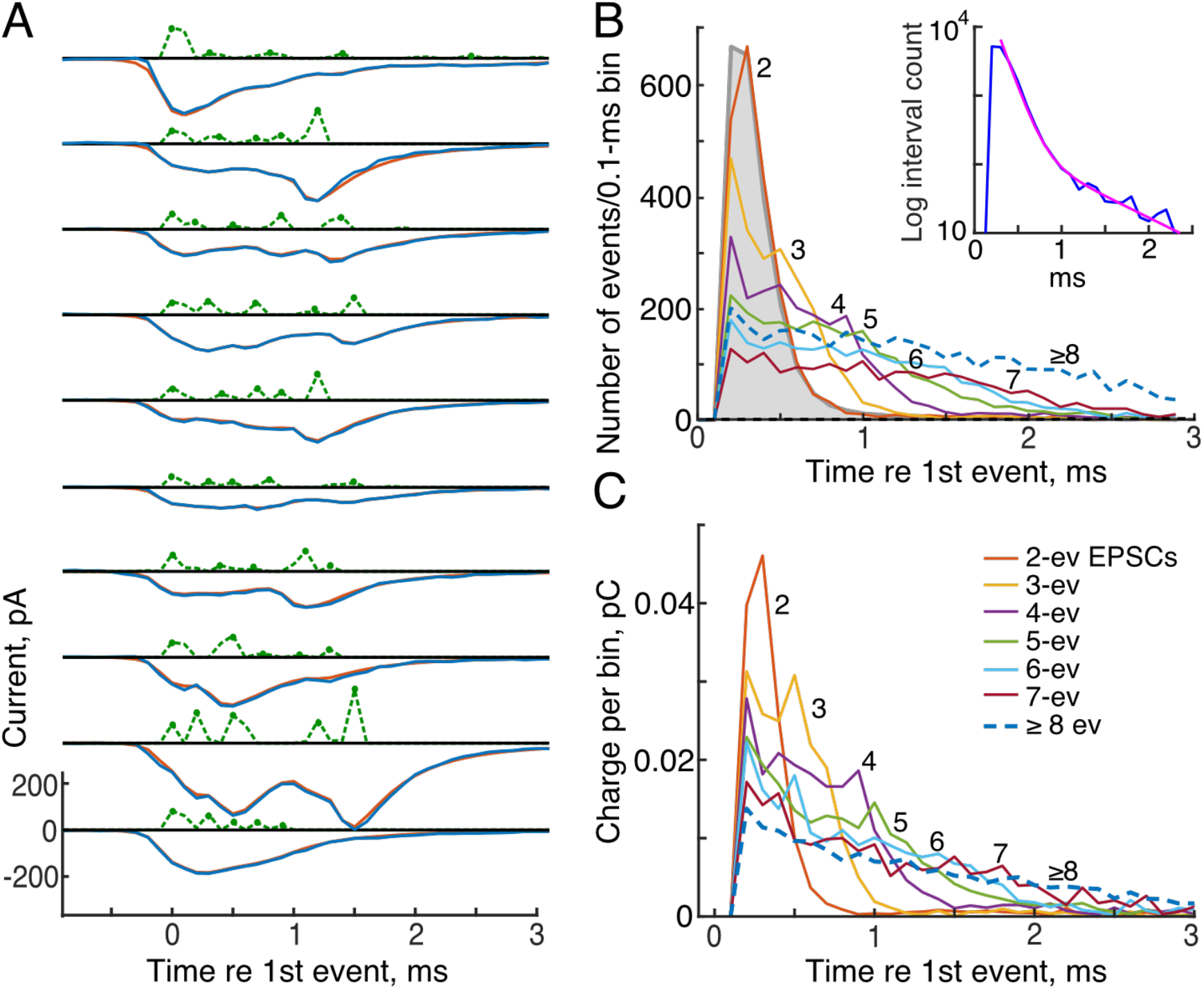
The evolution of release events through an EPSC. A. Waveforms of EPSCs (blue negative signals), kernel amplitudes (green dashed positive), and model fits (red) for ten examples of EPSCs with 5 events (green dots) from experiment T, plotted as in Figure 1C. B. Histograms of event occurrence times relative to the first event in the EPSC at time 0. The first events in the EPSC are off-scale and are not plotted. Colors identify data for EPSCs with different numbers of events,(legend in C and the numbers on the plots). Data from EPSCs with ≥8 events are combined into a single plot (blue dashed line). The interevent interval histogram for all events in the record is shown in filled gray. It is essentially the same as the 2-event histogram. The inset shows the interevent histogram on a logarithmic ordinate, with a best-fitting sum of two exponentials (time constants 0.13 and 0.74 ms, magenta). C. Like B, except plots of the charge injected versus time after the first event in the EPSC, for EPSCs with various numbers of events. First-event charges again are not shown.

As expected, the events are spread out over the duration of the EPSC. Figure 5B shows histograms of the event times averaged across all the EPSCs in experiment T containing two or more events. Time zero is the time of the first event in the EPSC, but the numbers of first events are not shown because they are perfectly aligned and so off-scale. In 2-event EPSCs (red-orange histogram labelled “2”), the second events occur centered around a single discrete maximum near 0.35 ms. In EPSCs with more events, indicated by the color code (legend in Figure 5C and numbers next to the curves), the events are spaced out over longer times; the total duration of the EPSC grows with the number of events. This result is complementary to that shown in Figure 2C: as the number of events increases, they spread out in time, leading to less summation of events and smaller EPSC amplitudes.

A histogram of all the interevent intervals (regardless of event number) from experiment T is shown by the gray histogram in Figure 5B. The histogram rises rapidly from zero to a mode near 0.3 ms and then decays with time, from 90% to 10%, over 0.34 ms (median 0.34 ms across 8 experiments). The decay is multi-exponential, shown on a logarithmic ordinate in the inset. A two-exponential fit to the decay (magenta line) has time constants 0.13 and 0.74 ms. The exponential decay shown in the inset is consistent with a random occurrence of events and not with a regular, clock-like occurrence. There is an apparent refractory period following the first event, in that the gray histogram has zero events in the bin at 0.1 ms, but any refractory period is very short in duration, one time-bin or a fraction of a bin, and is too short to be measured at the time resolution used here.

Figure 5B shows only the times of occurrence of events, not the amplitudes. A more useful measure of EPSCs is the amount of charge injected into the post-synapse at various times. This is shown in Figure 5C which plots the average profile of current injection in experiment T obtained by summing the event waveforms (like the green plots in Figure 5A), across all EPSCs with a given number of events, and converting the result to charge in the 0.1 ms bins of this histogram. The charge delivered by individual events is computed as the product of the event amplitude (current), the sampling increment, and the dimensionless area of the kernel (see Methods). Again, time zero is the time of the first event. The summed currents in the first events are again off-scale and not plotted. The charge curves decrease significantly as the number of events in the EPSC increases, as they must, given the constant area result of Figure 3.

The charges delivered by events in EPSCs with different event counts are shown in Figure 6; this figure shows averages (with SDs) of the charge injected by each event plotted against the sequence number of the event. Averages are shown for experiments T (Figure 6A) and H (Figure 6B) along with the average across all 8 experiments (Figure 6C). Each plot shows data from EPSCs with different numbers of events (legend in B). Consistent with the variability illustrated in Figure 5A, the SDs of each point are large, generally about equal to the mean.

**Figure 6.**
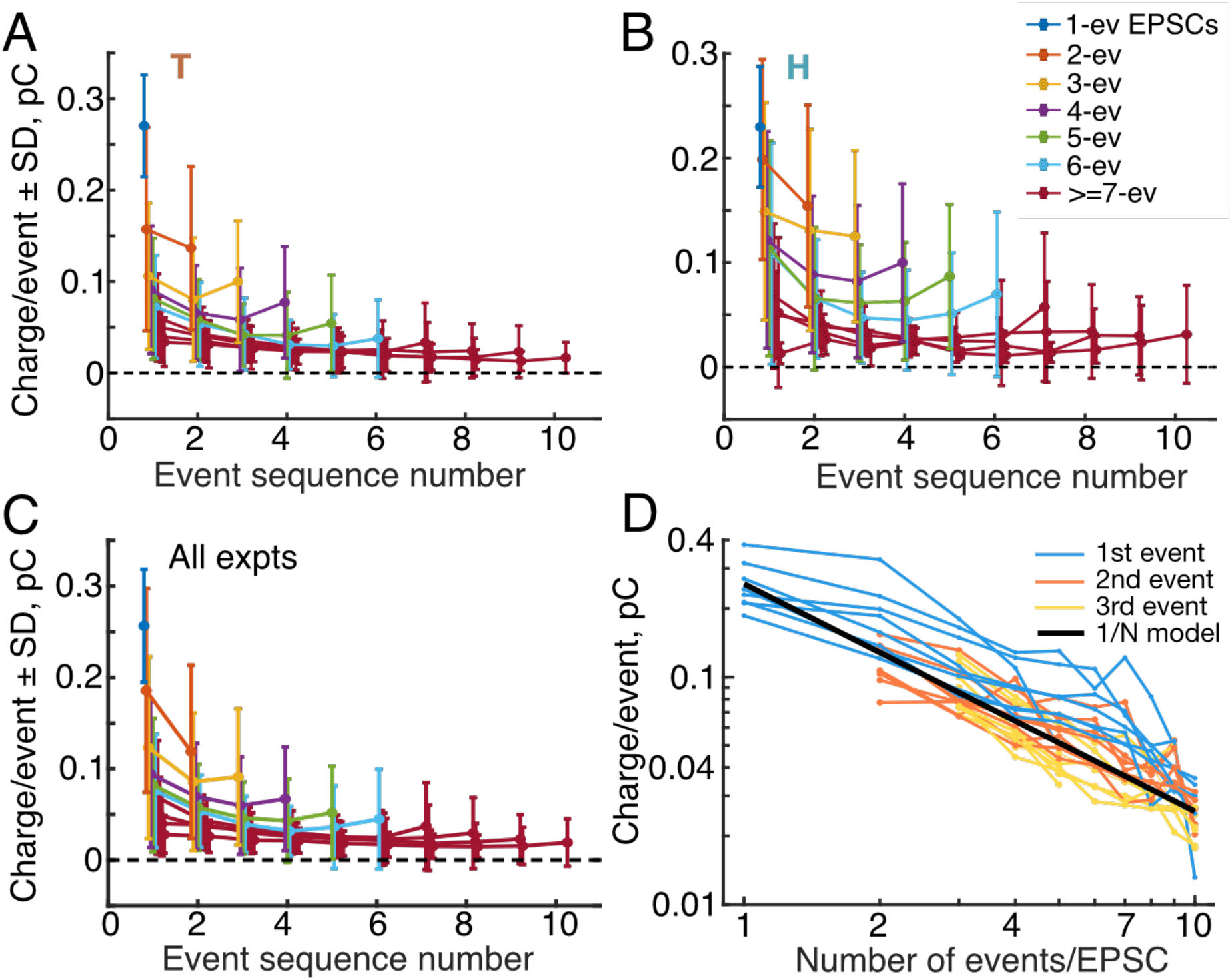
A. B. Mean and SD of charges delivered by individual events for EPSCs with various numbers of events (legend in B). The abscissa is the sequence number of the event (1st, 2nd, etc.), regardless of timing. Charges are computed as the sum of the amplitudes of the kernels in each event, converted to charge as described in Methods and normalized to charge/event. The data are from experiments T and H. The plots for ≥7 event EPSCs are shown in the same color because they overlap extensively and show similar behavior. C. Average of charge/event plots across all 8 experiments, with SDs. In A,B,C Points are displaced slightly along the abscissa to reduce overlap. D. Charge in the first, second, and third events (legend) of an EPSC versus the number of events. These data are the same as the first three points in each plot in parts A, B, C, for all 8 experiments. The black line shows the expected behavior if the charge in a monophasic EPSC is evenly divided among events in multiphasic EPSCs.. Note logarithmic axis scaling.

The average charge shows a shallow U shape through the EPSC, declining for early events and increasing for events near the end of the EPSC. That is, the first and last events tend to be larger than the events in between. More importantly, the charges get smaller as the number of events in the EPSC increases. Note that as the number of events increases, the decrease in charge is already apparent on the first event, on the first data point at left in each plot. This suggests that the amplitude of the boluses of neurotransmitter released in an EPSC is somehow determined at the beginning of the EPSC, as if the neurotransmitter is divided into smaller packets, which are then released in sequence.

The size of the smaller packets is hard to see in Figures 6A, 6B, and 6C. Figure 6D shows (blue lines) the charges of the 1st events in EPSCs plotted against the number of events in the EPSC; each line is from one of the 8 experiments. These are the first data points in parts A, B, and C of this figure. A simple behavior would be for the neurotransmitter to be divided into equal packets, so that the charges in the first events would decrease as 1/N, for EPSCs with N events. This idealized behavior is shown by the heavy black line which is (total charge)/N. For this line the “total charge” is the mean charge injected by monophasic (1-event) EPSCs. The blue lines are above the black line for EPSCs with 2 or more events. This means that a larger amount of neurotransmitter is released in the first event of a multiphasic EPSC than is predicted by equipartition of the monophasic EPSC.

For comparison, the orange and yellow lines show the same analysis for the 2nd and 3rd events in the EPSC. These are close to the inverse-N model, suggesting that equipartition holds for these events. As can be seen in Figures 6A and 6B, later events are smaller than the equipartition line, until the last one or two events in each sequence are larger again (not shown in part D).

## DISCUSSION

### The analysis

The results show that EPSCs in rat ANF dendrites can be resolved into a sum of individual release events consisting of current waveforms (kernels) having the same wave-shape as monophasic EPSCs. We assume that the release events correspond to pulsatile releases of boluses of neurotransmitter. The waveform of the kernel is determined by the cumulative transformation between the release of a pulse of neurotransmitter and the current in the afferent dendrite. It includes the diffusion of transmitter in the cleft, receptor binding, and receptor gating kinetics. This decomposition allows EPSCs to be classified by the number and asynchrony of the presumed release events making them up, and thereby provides a quantitative measure of monophasic and multiphasic EPSCs. The data show that multiphasic EPSCs are a minority fraction of the total, but may account for as many as 50% of the EPSCs in some preparations (Figure 2, Table 1).

The analysis assumes that the kernels summate linearly to produce an EPSC, requiring the assumption that the receptors are not saturated. Evidence of saturation was not observed (see Methods): the model fits uniformly well across the range of EPSC amplitudes; moreover the time constant of EPSC decay does not change with amplitude (Methods, Figure 7B). Thus the changes in waveform expected with saturation of responses to large neurotransmitter release were not observed. In principle, saturation can occur also for small events (Goutman and Glowatzki 2007, Figure 3). Low-level saturation was not seen, although this conclusion is less certain because of noise in small EPSCs.

**Figure 7.**
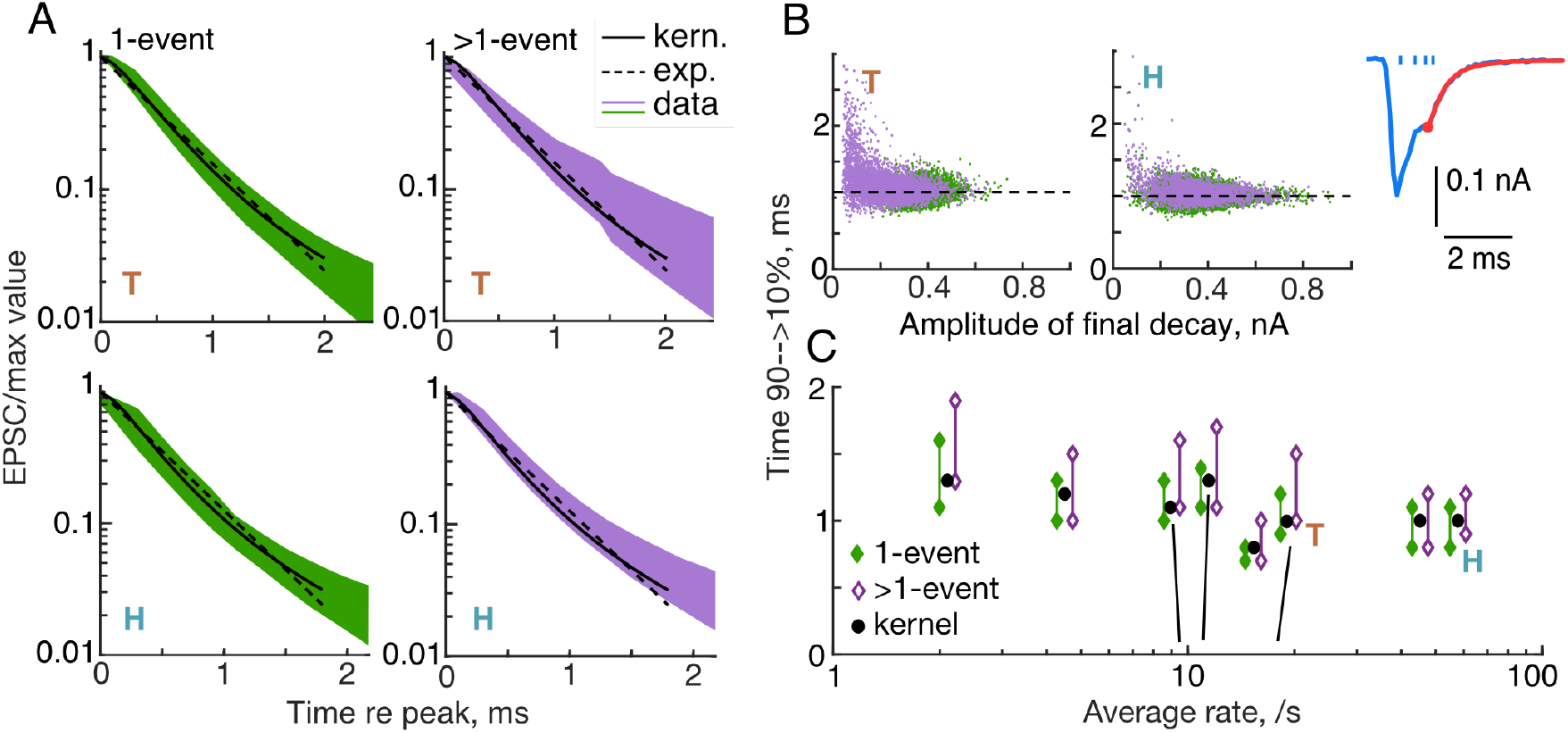
The decay time of the EPSC varies little with activity. Times for the EPSC to decay from 90% to 10% of its peak value were estimated from the final decay in the EPSC, after the last release event (e.g. for the red portion of the EPSC in the inset at upper right, starting at the red circle; the blue waveform is the whole EPSC and the ticks show the calculated event times). A. Final decays are plotted from experiments T (upper) and H (lower), separately for monophasic (1-event) EPSCs (left column, green) and multiphasic (>1-event) EPSCs (right column, purple). The waveforms are normalized to the beginning of the final decay and inverted for plotting, so they start at +1 at time 0. Solid black lines show the decay of the kernel and the green and purple bands contain the decays of all the waveforms between the 10th and 90th percentiles of the population of EPSCs at each time. Note that the >1 event EPSCs have roughly the same decay times as the 1-event decays. The dashed lines show the best single-exponential fits to the kernel decay. B. Final decay times versus event amplitudes for all the EPSCs in experiments T and H. Event amplitudes are the EPSC current at the start of the final decay (the red dot in the inset). The decay times are the same for the 1-event (green) and >1-event (purple) EPSCs, except when the final decay amplitude is very small, in which case the waveforms are corrupted by baseline noise. The horizontal dashed lines are the decay times of the kernels (1.08 and 0.97 ms). C. Summary of the decay times for eight experiments, plotted against the average rate of EPSCs. Each group of points shows the kernel decay time (black circle) and the 10th and 90th percentile range of the decay times for 1-event and >1-event EPSCs. For clarity, the data for experiments with rates between 8 and 20 /s are separated slightly along the x-axis. The correct rates are indicated by the black near-vertical lines.

### Are the EPSCs multiquantal?

In this analysis, multiphasic EPSCs result from the rapid sequential occurrence of individual smaller release events, whereas monophasic EPSCs result from a single synchronous release. This behavior raises the question of whether the release is multiquantal, representing synchronous (monophasic) or asynchronous (multiphasic) release of multiple vesicles (Glowatzki and Fuchs 2002; Keen and Hudspeth 2006; Li et al. 2000) or uniquantal, resulting from a single vesicle. With uniquantal release, multiphasic EPSCs would result from flickering of the fusion pore, producing multiple partial releases of the vesicle’s contents (Chapochnikov et al. 2014; Grabner and Moser 2018).

In the present data, the most important observation is that the releases generating an EPSC are made from a constant (except for the effects of noise) pool of neurotransmitter. This conclusion is suggested by two features of the data: (1) multiphasic EPSCs are smaller in amplitude than monophasic EPSCs, both individual events (Figures 6A, B, C) and the overall EPSCs (Figure 2A, D). This behavior is consistent with sharing a fixed pool of neurotransmitter by dividing it into smaller aliquots. (2) The total charge delivered by an EPSC, as measured by its area, is roughly constant, decreasing slightly as the number of events increases (Figure 3D; see also Grant et al. 2010; Chapochnikov et al. 2014). The area (charge) of an EPSC is linearly related to the amount of neurotransmitter released, as judged by pre-synaptic capacitance measurements (Li et al. 2009). Thus, within a single recording, EPSCs of all types seem to result, on average, from the same total amount of neurotransmitter release; by this is meant that the amount of neurotransmitter is, on average, the same and does not increase as the number of events in the EPSC increases. A constant neurotransmitter release could come about either from uniquantal release or from multiquantal release in which the average number of quanta released per EPSP is fixed.

Uniquantal release as proposed by Chapochnikov et al. (2014) seems consistent with most of the data in this paper. This mechanism assumes some variability in the fusion pore to explain the occurrence of mono- and multiphasic EPSCs. The randomness of the presumed flickers (Figures 5A and 5B) suggests a process in which open fusion pores are unstable, so that multiphasic EPSCs are produced by a variable and random sequence of openings. Recently the stability of fusion pores in CNS synapses has been shown to depend on the number of SNARE proteins recruited for the pore (Wu et al. 2017; Shi et al. 2012); a similar mechanism might apply to the IHC pore. Flickering might also result from a negative feedback interaction in the presynaptic release, such as inactivation of the presynaptic Ca channels when a fusion pore opens (Vincent et al. 2018).

Whatever the mechanism of flickering, it must be flexible enough to account for the range of partitions of the vesicle contents observed in the data (Figure 6), from the entire vesicle for a monophasic EPSC, to a number of partitions ranging up to a value of 10 or more in these data. An important property of that partition is that it occurs on the first opening and is maintained (except for random fluctuation) throughout the EPSC (see also Chapochnikov et al 2014, Figure 4E). It is not clear, for example, whether a mechanism like variability in the SNAREs recruited to form the fusion pore could produce the multiple possible partition sizes.

An additional question about uniquantal release is how the serial-dependence properties of EPSCs (Figure 4) arise. The phenomena in Figures 4B and 4C have time constants similar to those of synaptic depression (Goutman and Glowatzki 2007, 2011; Goutman 2012; Cho et al 2011) and the effect shown in Figure 4C represents a weakening of synaptic transmission, similar to synaptic depression. Depression seems to be mainly due to synaptic-vesicle depletion in this synapse, although there may be a small contribution of receptor desensitization (Goutman 2012, 2017). Anatomical studies show the presence of ~10 vesicles docked at the membrane under the synaptic bar (Khimich et al 2005; Chakrabarti et al. 2018), apparently in position for release. Assuming that these represent independent release sites with a low enough probability of release that usually only one site releases at a time (for a uniquantal EPSC), then the serial dependence of release asynchrony (Figure 4B) and the adaptation of EPSC charge (Figure 4C) are hard to explain. Presumably, parallel release sites should release independently and thus should not show serial dependence. That is, the only release site that should show the effects in Figures 4B and 4C is the one site that released on the previous EPSC. Given that there are multiple apparent release sites in a synaptic bar, it is not clear that the same site would release again and again, which seems to be necessary for uniquantal release and the effects in Figures 4B and 4C.

Multiquantal release is also consistent with much of the data, as long as the release is highly coordinated across release sites, so that the number of sites releasing is constant from EPSC to EPSC. Then small asynchrony in release times could produce multiphasic EPSCs with roughly constant area. Such a release model requires some mechanism, like cooperativity among release sites, to synchronize the releases and generate the constant release number. However, there must be enough randomness in the release times that multiphasic EPSCs are possible, i.e. there must be enough randomness to account for patterns like those in Figure 5A. These ideas were explored in a preliminary way with a hair cell synaptic model similar to that used by Peterson et al (2014) which contains multiple parallel release sites driven by a common signal, such as calcium. The synchronization of release across sites was produced by adding to the model strong cooperativity, in which release from one site increases the probability of release at other sites. With a delicate adjustment in the cooperativity strength, it is possible to achieve enough randomness in the release to produce results similar to Figures 5 and 6 but not so much that the number of sites releasing on a given EPSC varies, in which case the constant-EPSC-area constraint is violated. This fussiness of the model makes its results unconvincing.

It is useful to compare the present data on the rat hair-cell synapse with the properties of EPSCs in other ribbon synapses where the transmitter release is clearly multiquantal, e.g. ANF dendrites in the turtle (Schnee et al. 2013) and frog (Keen and Hudspeth 2006; Andor-Ardo et al., 2012; Li et al. 2009, 2014) and the retinal bipolar cell (Singer et al. 2004). The present data differs from these in two important ways. First, multiquantal-release synapses have mini-EPSCs that correspond to uniquantal release events. These are monophasic and are uniform in size and shape. In the usual theory used to describe EPSPs, mini-EPSCs serve the role of the kernel in the present analysis: larger EPSCs are constructed by increasing the number of vesicles released, thereby summing the mini-EPSCs. However there is no convincing evidence for mini-EPSCs in the present data set. In particular, there are very few monophasic EPSCs with small amplitudes (Figures 2A,B, Figure 2 - figure supplement 1, blue histograms) or areas (Figure 3A,B, Figure 3 - figure supplement 1) that would be suitable candidates for uniquantal mini-EPSCs. Indeed, the smallest amplitude EPSCs are multiphasic in the data analyzed here (e.g. the histogram for ≥4 event EPSCs in Figure 2 and Figure 2 - figure supplement 1), clearly inappropriate for a mini-EPSC.

The conclusion in the previous paragraph is tentative, because it may be necessary to reduce probability of release by a manipulation that lowers the calcium flux into IHCs in order to isolate mini-EPSCs (Li et al. 2009; Huang and Moser, 2018). That manipulation was not done in these data, in which the hair cells were depolarized with high-potassium solutions to achieve relatively high spontaneous EPSC rates. If recordings are performed in IHCs at fairly negative membrane potentials (~−70 mV; Niwa et al, 2020, personal communication, Figure 3), calcium influx into IHCs is small, shown by a lack of spontaneous firing. In this condition, compared to recordings with more depolarized IHCs, EPSCs tend to have smaller amplitudes and area as well as a low number of events per EPSC, and some of these small and fast events have the expected shape of mini-EPSCs.

The second difference between the present data and the multiquantal EPSCs of Li et al. (2009) is in the histograms of EPSC amplitudes (Figure 2) or areas (Figure 3). For monophasic EPSCs, these are symmetrical and narrow (median CV=0.20 for amplitudes and 0.25 for area over 8 experiments), comparable to uniquantal EPSC amplitudes in frogs (Li et al. 2009). For multiphasic EPSCs, there is a substantial tail in the amplitude distribution (but not the area distribution) on the low-amplitude side (Figure 2A and Figure 2 - figure supplement 1, see also Grant et al 2010). By contrast, multiquantal amplitude histograms have a substantial tail on the large-amplitude side of the amplitude and area distributions (Singer et al. 2004; Li et al. 2009; Andor-Ardo et al. 2012). This large-amplitude multiquantal tail, of course results from the mixture of EPSCs containing different numbers of quanta. Similar tails are observed here in EPSC area distributions, but only with overlapping of monophasic EPSCs in high-rate data sets like Experiment H (Figure 3B), as discussed previously.

There is a small decline in the area of EPSCs with larger numbers of release events (Figure 3D); this is a robust phenomenon, observed in all experiments. It seems to reflect some loss of neurotransmitter through a multi-event release. Chapochnikov and colleagues (2014) argued against receptor desensitization as a mechanism for these declines, using modeling. Receptor desensitization also seems inconsistent with (1) the randomness of event sizes and the occurrence of larger release events near the end of multiphasic EPSCs as in some of the examples in Figure 5A; (2) the upturn in average release event size near the end of multiphasic EPSCs in Figure 6A,B,C; that is, there is no evidence of a monotonic decline of event size during the EPSC, as would be produced by desensitization. Instead one is led to consider mechanisms like failure to release some fraction of the neurotransmitter during an extended release (as occurs in the “kiss-and-run” phenomenon in other synapses; Allabi and Tsien, 2014), loss of neurotransmitter through an extended release sequence, a predilection for smaller pools of neurotransmitter to be released asynchronously, or restricted permeation through a uniquantal release pore in multiphasic releases.

### Quantal sizes

The presynaptic quantal size of an EPSC can be estimated with simultaneous presynaptic capacitance recording and postsynaptic EPSC recording. With this method, Li and collaborators (Li et al 2009) estimated the postsynaptic EPSC charge injection from a single presynaptic vesicle to be about 45 fC, as in mini-EPSCs in the frog hair-cell synapse. This corresponds reasonably well to the charge injected by the smallest events in the data of Figure 6, roughly 10-30 fC. It is much smaller than the charge required to produce monophasic EPSC in Figure 3, which is 200-300 fC.

The data in the previous paragraph argue for a multiquantal mechanism in the rat hair cell. The experiment of Li and collaborators has not been done in a mammal; however capacitance recordings in hair cells have been done (e.g. Grabner and Moser 2018; Neef et al. 2007), in which it was possible to resolve single steps in the capacitance data that seem to correspond to single vesicle releases. These capacitance steps had the amplitudes expected for one or two single synaptic vesicles, based on synaptic vesicle surface area. However, postsynaptic recordings were not made in those experiments so the amount of charge injected by the resulting EPSCs was not determined. Thus the question of quantal size requires further research.

### Final comments

In our opinion, the question of multiquantal versus uniquantal release remains open. The uniquantal flickering-release model is supported by the near-constancy of EPSC charge and the lack of a convincing mini-EPSC in the present data. However experimental verification that a single vesicle can produce the large charge injections required for the postsynaptic EPSCs in these datasets is missing. Multiquantal release is also consistent with much of the data, however, one has to assume that release is highly coordinated across release sites, to support near-constancy of release charge, with enough variability to produce both monophasic and multiphasic EPSCs.

The flickering model is similar to the “kiss-and-run” phenomenon which has been proposed for CNS synapses (Alabi and Tsien, 2013; He et al 2006). In both, the fusion pore closes after a partial release of its contents. In the case of kiss-and-run, there is only a single opening of the pore, after which the vesicle is recycled in a way that speeds its availability for another release.The evidence presented here and by Chapochnikov et al (2014) for uniquantal flickering suggests a different process in which the pore repeatedly opens and closes until the vesicle contents are released. These differences in function could derive from the different molecular constitution of the vesicle fusion pore in hair cells, relative to CNS synapses (Nouvain et al 2011).

## MATERIALS AND METHODS

### Experimental methods

All aspects of the animal preparation were approved by the Johns Hopkins Animal Care and Use Committee. The data used here overlap with data previously reported (Wu et al 2016), some of which were obtained in the same experimental preparations. Briefly, Sprague-Dawley rats of either sex and ages between postnatal day (P) 17 to 31 were deeply anesthetized with isoflurane and decapitated. The cochleae were removed and the apical cochlear coil placed in a recording chamber in extracellular solution (in mM: 5.8 KCl, 144 NaCl, 0.9 MgCl2, 1.3 CaCl2, 0.7 NaH2PO4, 5.6 glucose, 10 HEPES, pH adjusted to 7.4 with NaOH, at 300 mOsm). The bath temperature was 22-25°C. The preparation was viewed using differential interference contrast optics which allowed ANF dendritic terminals on IHCs to be visualized. Intracellular recordings from single dendritic terminals were obtained by whole-cell patch using pipettes of resistance 11-15 MΩ; the series resistance was <60 MΩ and was not compensated. The cells were voltage-clamped (pCLAMP 9.2 software and a Multiclamp 700A amplifier, Molecular Devices) at membrane potentials between −99 and −84 mV. Membrane current was digitized at 50 kHz sampling rate, low-pass filtered at 10 kHz, and recorded for further analysis. The data consist of long runs (up to 600 s) of EPSCs recorded in 8 experiments.

There was no explicit stimulation of the hair cells, except that they were depolarized with an elevated potassium concentration in the bath solution (5.8 mM in 3 cases, 15 mM in 5 cases), with corresponding decreases in sodium concentration. To check the IHC potentials, intracellular recordings were performed from 10 IHCs at P24: in 5.8 mM K^+^, the IHC membrane potential was near ‘rest’, at −60.4 ± 7.4 mV, and in 15 mM K^+^ it was depolarized to −40.5 ± 12.0 mV. The rate of EPSCs varied from 2 to 58 /s in different preparations (Table 1). There was no systematic difference between data at the two potassium concentrations (see also Grant et al. 2010); across the preparations, the median steady-state rates were 15.3 /s (5.8 mM) and 10.2 /s (15 mM), respectively (P=0.6, ranksum). Maintaining the HC potential near and above rest raises the calcium concentration in the IHCs and provides a roughly linear dependence of postsynaptic response on calcium entry (Goutman and Glowatzki 2007), more similar to in-vivo conditions than if the IHCs were not depolarized.

The data analyzed in this paper come from the 8 experiments summarized in Table 1. These were cases in which the experimental preparation and the properties of the EPSCs were stable through the entire recording period. The properties reported here were similar across all experiments, except as noted in Results. There were fluctuations in rate during each recording, like those described previously in loose-patch recording (Wu et al. 2016, Figure 2) and *in-vivo* extracellular recording (Teich et al. 1990). These were not accompanied by changes in EPSC amplitude or other characteristics. Note that rate was not systematically controlled or evaluated, e.g. with a time-varying stimulus. Data from five additional experiments were analyzed but not included here. In three of these experiments, substantial changes in holding current and the properties of EPSCs occurred during the recording time, suggesting damage to the preparation. The data in two other experiments showed unusually high numbers of multiphasic EPSCs (>50%), which had properties that were similar to the two damaged preparations mentioned above. All five excluded data sets also showed substantial differences between the time constants of monophasic and multiphasic EPSCs in terms of decay time constant (as explained in Figure 7), not typical of the 8 included experiments. This behavior suggests substantial nonlinearity in the EPSCs, which means that the assumptions necessary for the analysis do not hold for these experiments.

### Analysis

EPSC recordings were downsampled from 50 kHz to 10 kHz using the Matlab resample() function. This function first low-pass filters the data with a sharp antialiasing filter (cutoff frequency 5 kHz) and then resamples. The filtering and resampling reduce noise in the data record (by about 10 dB), by eliminating high-frequency (noise) stimulus components. This manipulation does not change the wave shapes or amplitudes of the EPSCs, which have low-pass spectra with a cutoff frequency of ~2.5 kHz. Reducing the sampling rate is also convenient because it reduces the time required for the model-fitting calculations described below. Some experiments were re-analyzed using 20 kHz resampling, with no change in the qualitative results.

EPSCs occur as isolated signals separated by periods of baseline (examples in Figure 1A). Because spontaneous EPSCs occur randomly in time, on some occasions EPSCs overlap partially; at the rates observed here, overlap was infrequent (11% of EPSCs for the highest EPSC rate, 58 /s, less for the others). No attempt to separate overlapping EPSCs was made, and the overlapped pairs were analyzed as one EPSC (discussed further in connection with Figure 3). In the raw data, the baseline between EPSCs was equal to the holding current and showed slow variations through a recording. To set the baseline to zero, the portions of the data record remaining after excluding EPSCs were fit with a smooth function computed by averaging non-EPSC points in overlapping 10 ms windows and interpolating (linearly) to the sampling rate. The resulting interpolated function was an estimate of the baseline and was subtracted from the data record, including during the EPSCs, to set the baseline to zero. Note that this method does not modify the EPSC waveforms. EPSCs were delimited, for baseline zeroing, by setting a threshold (typically −15 to −30 pA relative to the baseline), estimated by manually scrolling through the data set, looking for a threshold that did not exclude the smallest EPSCs. These thresholds are not limited by the baseline noise; the ratio of the EPSC threshold to the rms value of the baseline noise is 9.3 to 15.7 in 8 experiments, median 13.6. EPSCs were then defined as compact regions where current was more negative than the threshold. EPSC definition was done once crudely to allow baseline zeroing and then again after baseline correction. The black bars in Figure 1A show the EPSC extents resulting from thresholding. To ensure that small EPSCs were not missed, the waveforms of the largest currents remaining in the data record after EPSC detection and removal were displayed; if these waveforms seemed to be EPSCs based on their waveforms and durations, the threshold for EPSC detection was adjusted. The number of such waveforms was small (less than 100 in data sets with thousands of EPSCs).

To analyze the EPSCs, each was approximated as a sum of elementary waveforms, called the *kernel* below. The majority of EPSCs are monophasic (Table 1) and these monophasic EPSCs have very similar waveforms, with different amplitudes. We assumed that the waveform of monophasic EPSCs is the kernel, which was estimated by averaging monophasic EPSCs, after normalizing them to an amplitude of −1. EPSCs were included in the average only if they were preceded and followed for 10 ms by baseline, i.e. no nearby EPSCs, and not part of an overlapping pair of EPSCs. Multiphasic waveforms were manually edited out of the average by superimposing all apparently monophasic waveforms (normalized to the same maximum amplitude and aligned on their peaks) in an interactive display, and then setting limits on the waveforms included. Most (median 89%) of the waveforms ultimately determined to be monophasic in the data were included in the averages, which were only slightly different in different experiments, mainly in the rise-fall time (last column of Table 1; Figure 1 - figure supplement 1). In terms of this averaged kernel waveform, *k(t)*, an EPSC can be approximated by the following:

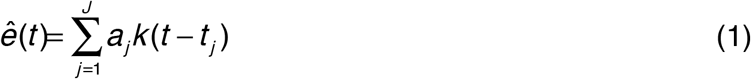

*ê(t)* is the sum of *J* kernels with amplitudes *a_j_* and occurrence times *t_j_* (actually the time of the minimum of the kernel). An example of such a sum is shown in Figure 1B where the EPSC (blue negative) is fit by the red dashed waveform, which is the sum of the green kernels. The lines in the top half of Figure 1B show the amplitude *a_j_* of each kernel; they are located at the kernel’s time delay *t_j_*. Each EPSC *e(t)* was fit individually by such a sum by choosing *J* and the amplitudes {*a_j_*} and times {*t_j_*} so as to minimize the error between the EPSC waveform and *ê(t).* The error is given by

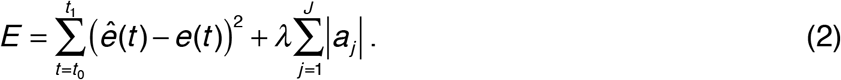

The first term in *E* is the squared difference between the EPSC *e(t)* and the fit function *ê(t)* from Eqn. 1. The difference is computed over a time range [*t_0_, t_1_*] which is the extent of the EPSP determined by thresholding (the black bars in Figure 1A), extended by 2 ms in both directions. The baseline extension is done to include baseline adjacent to the EPSC, which provides a zero reference that forces the DC term in the fit to 0, consistent with Eqn. (1) which does not have a constant term. Because the data are noisy, the squared difference component of *E* can always be minimized by increasing *J,* the number of kernels, up to the limit of one kernel in each sampling time increment, which causes addition of small kernels that improve the fit. However, as more smaller kernels are added, they are likely fitting noise in the data. The second term in the error is designed to avoid this by penalizing the addition of more kernels. This method, called lasso (Tibshirani, 1996), allows the addition of another kernel to the sum (increasing *J* and adding another *a_j_*-sized kernel at time *t_j_*) only if it reduces the squared error (first term in Eqn. 2) more than it increases the penalty, equal to the sum of the kernel amplitudes multiplied by λ (second term in Eqn. 2). The penalty term increases as more kernels are added.

The parameter λ sets the relative importance of the squared-difference and the penalty. Increasing λ reduces the number of kernels at the cost of an increase in squared error. The Matlab function lasso() was used to fit each EPSC individually. λ was set to the value at which the squared error exceeds its minimum by 1 SEM (estimated by 5-fold cross-validation). As shown by the examples in Figure 1, the errors in the fits are small (but not as small as they could be with no penalty). The accuracy of the fit can be measured by the *relative root-mean-square error*, equal to the root-mean-square error divided by the root-mean-square EPSC value. The relative error was 0.0087 (median across experiments) for monophasic EPSCs (*J*=1) and 0.0034, 00033, and 0.0028 for EPSCs with *J* = 2, 3, and >3. The fact that the error for multi-component fits (*J*>1) is small supports the validity of the assumption in Eqn. 1. This calculation is a form of deconvolution; different approaches to this problem for the hair-cell/auditory-dendrite synapse are described by Andor-Ardo et al. (2012) and Chapochnikov et al. (2014).

In the body of the paper, the kernels making up a fit are called *events.* These events are assumed to correspond to releases of boluses of neurotransmitter. In order to classify EPSCs according to the number of presumed neurotransmitter releases they contain, events were counted, except that kernels located in adjacent time samples were combined into one event, under the assumption that they may represent a release event spanning the two bins. The results are called *events* in the body of the paper. Usually, events contain one kernel except when combined in this way, as for the 4th and 5th kernels in Figure 1C.Thus the EPSC in Figure 1C is classified as having 4 events, even though it contains 5 kernels. The event counts shown in Figures 1A and 1D are calculated in this way.

In many cases of EPSCs with 2 and 3 events, there is one large event and one or more very small events. An example is the 6th EPSC in Figure 1A, in which there is one very small event evident in the green trace. Because such small events may represent noise, the small event is not counted in classifying the EPSC, in cases where the areas of small events are less than 0.03 times the area of the largest event in the EPSC. Thus in the example in Figure 1A, the sixth EPSC is labelled as having one event. This reclassification was done only for EPSCs having two or three events. The results seem reasonable in terms of the shapes of the EPSCs, in that EPSCs with *J=2* kernels in the fit, with one very small one, usually have waveforms that are essentially the same as monophasic EPSCs. None of the results described here are dependent on this reclassification, except in minor quantitative details. To check its effect, the threshold was raised to 0.1. At this threshold the median fraction of EPSCs modified by removing a small event was 0.07 (range 0.03 to 0.12 in 8 experiments).

#### Amplitude, area, and statistics

The amplitudes of kernels *a_j_* have units pA. The kernel itself is always normalized to have maximum amplitude −1, so is dimensionless. The areas of EPSCs and events are expressed as charge. For EPSCs, this is computed straightforwardly as the sum of the EPSC current (the blue area in Figure 1C) multiplied by the sampling time increment (0.1 ms) yielding units of Coulombs. For events, the kernel amplitudes (*a_j_*) is multiplied by the sampling time increment and by the area of the dimensionless kernel (mean 7.10, SD 0.89, for 8 experiments). The result is the charge (C) delivered by that event. The charge of an event is the sum of its kernel charges. The sum of the event charges is approximately the same as the overall EPSC charge, as it must be from Eqn. 1.

Statistical tests on the data are described as they are used. Usually non-parametric methods were used (rank-sum) except that least-squares line-fitting and cross-correlation (Pearson) are used to evaluate trends in the data. Statistics were calculated with routines in the Matlab statistics toolbox: signrank(), ranksum(), corr(), polyfit().

### Linearity in the deconvolution of EPSCs

The analytical approach used here assumes that an EPSC is a linear sum of currents (Eqn. 1) produced by releases of boluses of neurotransmitter (the kernels). Because neurotransmitter receptor kinetics contains nonlinearities and non-stationarities (e.g. Destexe et al. 1995; Häusser and Roth 1997), in that rate constants depend on the concentration of neurotransmitter and the state of the receptor molecule, there is reason to question whether EPSC generation is as linear as assumed.

A necessary condition for non-stationarity, which may also detect nonlinearity, is based on the time constant of decay of the EPSCs. This time constant is sensitive to changes in the open-state rate constants of the receptor molecules. The simplest time constant to measure is that associated with the final decay of the EPSC, after the last release event. In the linear summation model, that decay should be the same as the decay of the kernel, because the EPSC decay is the sum of one or more decaying kernels (Figure 1B). An example of the final decay is shown by the red tail on the inset EPSC at upper right in Figure 7. Final decay is defined here as the decay of the EPSC after the last event.

The EPSC decays are remarkably similar. Figure 7A shows decay waveforms from two example experiments (called “T” and “H” in all figures). The waveforms are inverted and normalized by the current at the beginning of the final decay (e.g. the current at the red dot in the inset of Figure 7). The colored bands show the region occupied by all the waveforms between the 10th and the 90th percentiles of the population at each time point. The solid black lines show the decay of the kernel waveforms from these two experiments. The decays are plotted on a log ordinate, so a single exponential decay is a straight line. The dashed black lines show the exponential that fits best the decay of the kernel. Both the kernels and the raw data (colored bands) are seen to be multiexponential, because their waveforms are not straight lines in this plot. For 8 experiments the median values of the first two decay time constants are 0.50 ms and 1.47 ms. Because they are multiexponential (see also Häusser and Roth 1997), the decays are quantified in the rest of the figure as the time required to decay from 90% to 10% of the peak amplitudes (median value 1.0 ms across 8 experiments, Table 1). Using the kernel decays as a reference for comparison, it is clear that the decay times are very similar for monophasic EPSCs (left column of Figure 7A, green) and multiphasic EPSCs (right column, purple) for both experiments T and H.

The effect of current amplitude is shown in Figure 7B where 90-10 decay time is plotted against the amplitude of the EPSC at the final event (the current at the red dot in the inset) There is no effect of amplitude, except for very small amplitudes, where the decay times cover a broad range. However, at small amplitudes, the decay waveforms are strongly affected by baseline noise currents and it seems likely that the broadening of the time constant range represents noise in the measurement rather than a systematic effect.

The results for experiments T and H are typical of all eight experiments (Figure 7C) in that the kernel decay time is similar to the decay times from both monophasic and multiphasic EPSCs. There is a weak dependence of decay time on average EPSC rate (for the kernel decay times in Figure 7C: r=−0.65, P<0.09). This marginal effect may reflect variability of experimental conditions from experiment to experiment, such as temperature (Wu et al. 2016, Figure 3C).

The behavior shown in Figure 7 is a necessary condition for the linearity assumption to be valid. In the five experiments not included in this analysis, the decay times were different for monophasic and multiphasic EPSCs, suggesting the existence of nonlinearities in those experiments.

Despite the result in Figure 7B, there was a small tendency for the halfwidth of monophasic EPSCs to increase as the amplitude of the EPSCs increases. This broadening of EPSCs suggests a small nonlinearity in the receptor kinetics. However, given the constancy of the decay time constant in Figure 7B, it seems more likely that this effect is due to a slight asynchrony of the neurotransmitter boluses as the amount of transmitter increases.

## ACKNOWLEDGEMENTS

This work was supported by National Institute on Deafness and Other Communication Disorders Grants R01DC006476 and R01DC012957 to E.G. and National Institute on Deafness and Other Communication Disorders P30 DC005211 to the Center for Hearing and Balance at Johns Hopkins, the David M. Rubenstein Fund for Hearing Research and the John Mitchell Jr. Trust. We thank Eunyoung Yi and Lisa Grant for recordings that were included in the analysis of EPSCs and Philippe Vincent for measurements of hair-cell membrane potential.

---

**Figure 1-figure supplement 1.**
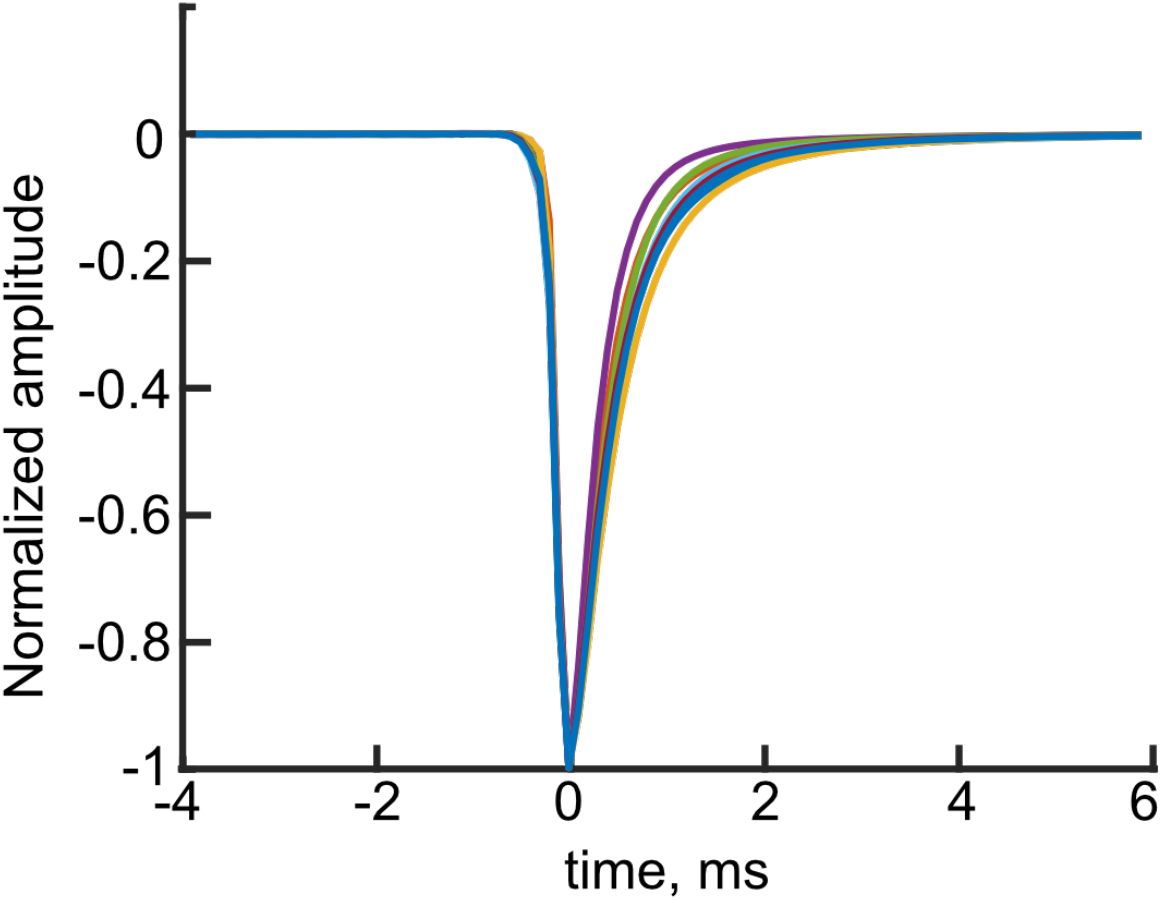
Kernel waveforms for the 8 experiments. Kernels derived by averaging monophasic EPSCs. These are *k(t)* in Eqn. 1.

**Figure 2-figure supplement 1.**
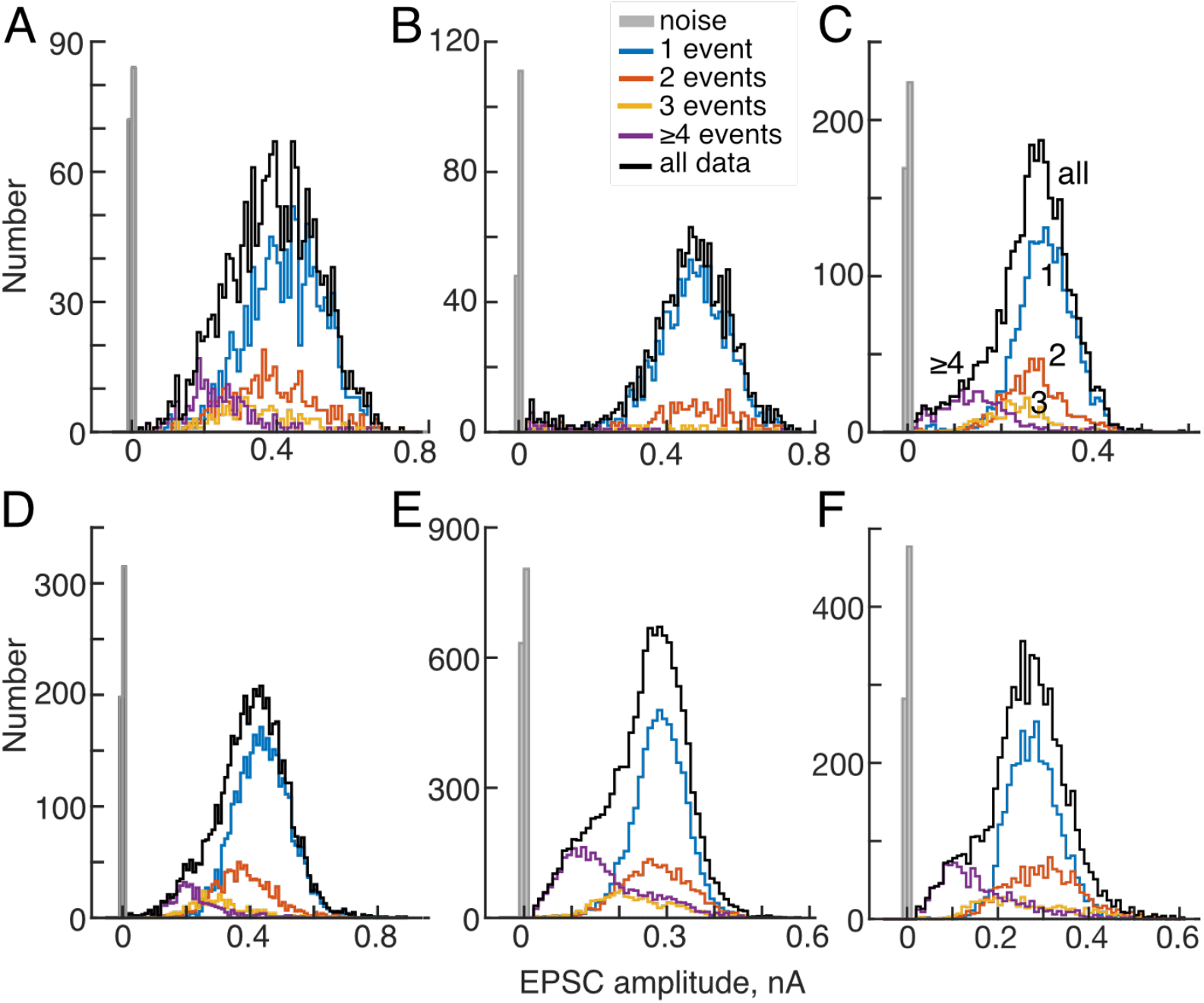
Amplitudes of EPSCs of six experiments not shown in Figure 2A,B. Histograms of EPSC amplitudes from six experiments, plotted as for Figure 2A, B. These are mostly similar to Figure 2A in that distributions move toward lower amplitudes for larger number of events, as labelled in the top center plot. In all cases, the low-amplitude EPSCs are mostly or entirely multiphasic waveforms (event numbers ≥4). The data in A and D were studied in 5.8 mM K^+^ and the remainder were studied in 15 mM K^+^.

**Figure 3-figure supplement 1.**
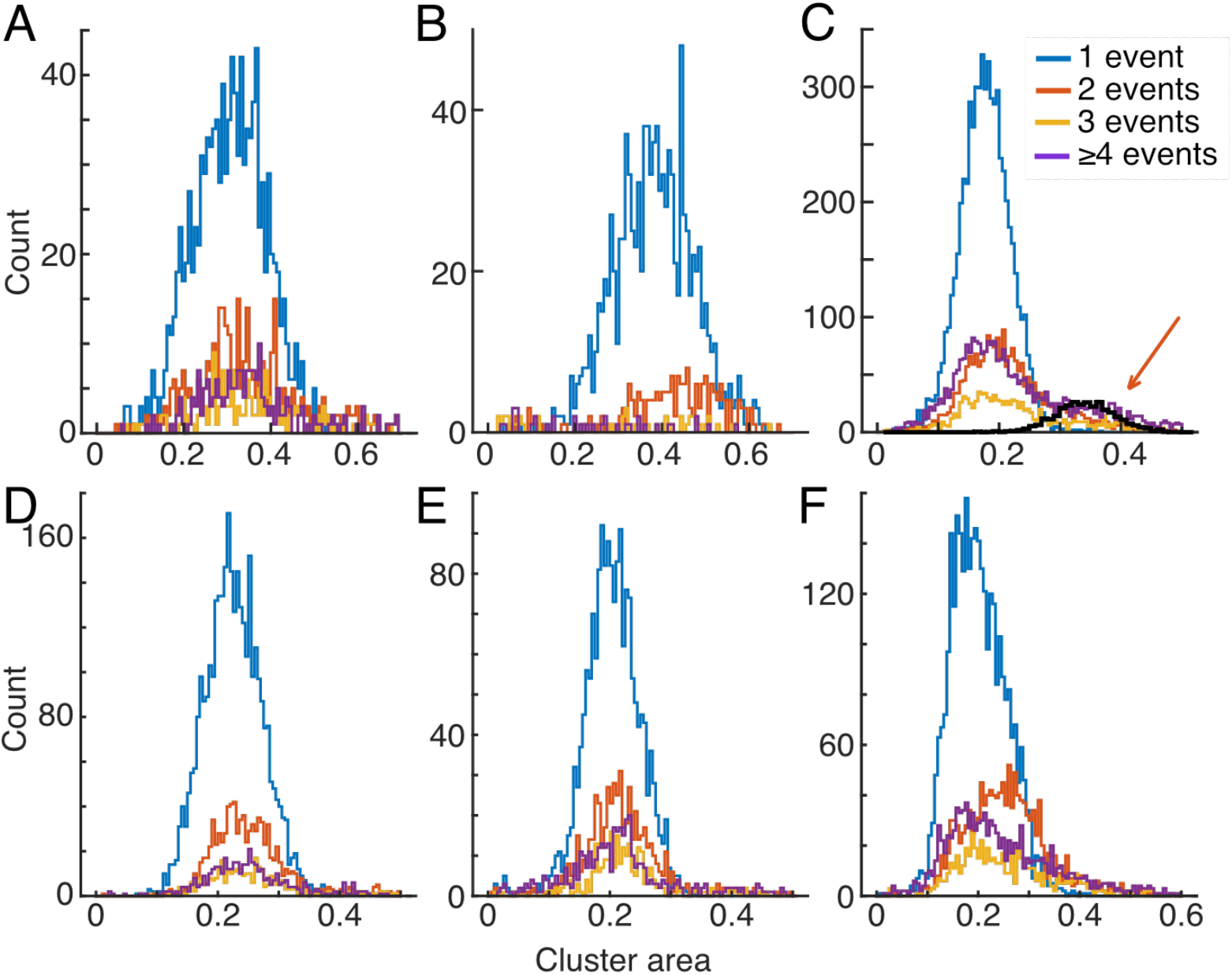
Distributions of EPSC area for the six experiments not shown in Figure 3 (arranged in the same order as Figure 2-supplement 1). The distributions for multievent EPSCs usually center around the same mean as the histogram for 1-event EPSCs, except for the experiments shown in B and F. C shows data from an experiment with high EPSC rate (45 /s); as for experiment H, there are secondary peaks in the distribution at the place predicted by the simulation of overlapping EPSCs (heavy black plot, orange arrow.

